# Anatomical and functional organization of the interpeduncular nucleus in larval zebrafish

**DOI:** 10.1101/2024.10.09.617353

**Authors:** You Kure Wu, Luigi Petrucco, Ruben Portugues

## Abstract

The habenulo-interpeduncular pathway is a highly conserved neural circuit across vertebrates, but the anatomical and functional architecture of the interpeduncular nucleus (IPN) remains poorly understood. Here, we use a combination of immunohistochemistry, volumetric electron microscopy (EM), and two-photon imaging to provide the first detailed characterization of the internal organization of the IPN in larval zebrafish. We show that the IPN receives extensive projections from the tegmentum, and reveal a strict segregation between the dorsal (dIPN) and ventral (vIPN) subcircuits, with minimal cross-communication. In the dIPN, we characterise in detail the inputs and outputs of r1π neurons, which have been recently identified as representing the animal’s heading direction. In the vIPN, we identify six distinct glomerular structures, each exhibiting specific patterns of reciprocal connections and projection pathways. Finally, we demonstrate that the connectivity and spontaneous activity patterns of habenular axons are shaped by the local anatomical features of the IPN, suggesting a role for the local interneurons in modulating presynaptic dynamics. Together, these results enhance our understanding of the internal organization of the IPN, and provide a framework for future investigations into both its physiology and its involvement in behavior.

## Introduction

The habenulo-interpeduncular circuit is a fundamental and highly conserved neural pathway across vertebrate species, including fish, amphibians, birds, and mammals [1–3]. This circuit is primarily defined by the fasciculus retroflexus (FR), a prominent fiber bundle that projects glutamatergic and cholinergic fibers [4] from the dorsal habenula (in teleost fishes, medial habenula in mammals) [5, 6], which is located bilaterally in the dorsal diencephalon to the interpeduncular nucleus (IPN). The IPN, situated in the ventral part of the rostral-most hindbrain, is a region of dense neuropil whose borders are defined by the projection territory of the habenular axons.

The morphology and physiology of local IPN neurons is still largely unknown. It is known that they are mostly GABAergic cells of varied morphologies. Early reports from Golgi preparations have described two types of interpeduncular neurons: one with complex dendritic trees, and another one with smaller and spiny dendrites; howeer a thorough classification is still missing [7–9]. These local IPN neurons receive inputs from the habenula and dorsal regions of the tegmentum [10–12], and project back to tegmental areas [12] and to more caudal hindbrain territories such as the griseum centrale in fish [13].

The IPN has been implicated in a range of neural processes. It is thought to play a role in linking limbic, sensory, and telencephalic systems to motor centers, thereby integrating emotional and cognitive information with motor functions [7]. In zebrafish, the left and the right habenula have been shown to be highly responsive to visual and olfactory stimuli, respectively [14, 15]. Ablation and optogenetic experiments have implicated the IPN in odor avoidance and fear responses [16, 17], place avoidance learning [18–22], familiarity learning [23], reward encoding and addiction [23–27], stress [28, 29], social behavior [30, 31] and, more recently, navigation. Indeed, reciprocal projections link the IPN with the tegmental areas where the most basal head-direction cells can be found in mammals [12, 32, 33], and ablation studies suggest the involvement of the IPN in allocentric [34–37] and egocentric navigation [38]. More recently, data from our lab have highlighted how this region could be crucially involved in organizing the activity of a ring attractor network that integrates heading direction [39, 40], and in mediating homeostatic navigation [41].

Despite its anatomical prominence and proposed functions, the IPN remains a poorly understood structure with a largely unknown internal organization and functional dynamics. It is located in a very ventral and relatively inaccessible part of the brain, which has made it challenging to study in vivo [5]. Its dense neuropil composition and the compact arrangement of its neuronal circuits further complicate the systematic description of its anatomy by classical staining and tracing studies. Overall, detailed insights into its internal physiology and anatomy remain scarce.

Here, we employed a combination of techniques including immunohistochemistry, targeted labeling of photoactivatable fluorescent proteins, volumetric electron microscopy (EM) reconstructions, and two-photon imaging. With these methods, we thoroughly characterize the internal structure of the IPN, its relationship with surrounding neural areas, and some of its dynamic properties. Through these approaches, we provide the first systematic description of the internal organization and anatomical relations of the teleost IPN.

We show that the IPN has extensive relationships with surrounding brain areas, and it exhibits a strict segregation between the dorsal and ventral sub-circuits. The dorsal part of the IPN is characterized by a continuous and mixed distribution of neurons, where adjacent arborizations partially overlap, while the ventral part displays a highly compartmentalized structure organized in six distinct glomeruli. Specific regions of the dorsal hindbrain contribute axons and dendrites to the IPN, and can be considered part of the IPN circuit. Additionally, the local dynamics of habenular axons within the IPN appears to be strongly shaped by the local anatomical structure. These findings significantly advance our understanding of the IPN’s internal architecture and its functional connectivity, providing a comprehensive framework for future investigations into its role in vertebrate neural circuits.

## Results

### Cytoarchitecture of the IPN in larval zebrafish

We started our investigation of the larval zebrafish IPN by performing detailed confocal reconstructions of the structure’s volume in different transgenic lines at 7 dpf, and registering the obtained data to a standard reference space, sampled at isometric 0.5 × 0.5 × 0.5 µm voxels. We have also considered for the analysis the volume of the IPN that can be observed in the EM dataset described in [42], for which we have found a transformation to the same standard reference space.

Boundaries of the region were defined looking at the region spanned by the left and right habenula projections Figure 1A. As previously reported, the zebrafish IPN volume consists mostly of a dense neuropil mesh that can be separated in a dorsal (dIPN) and a ventral (vIPN) part separated by a central core of densely packed somas. Analyzing the EM stack in this region reveals that at 5 dpf, there are approximately 130 somas in the core. We refer to neurons with the soma in the core as IPN core neurons. The core extends in stripes that join seamlessly other areas of densely packed somas, anteriorly in what we define “anterior wall”, and dorsally in the medial raphe (Figure 1B).

**Figure 1:**
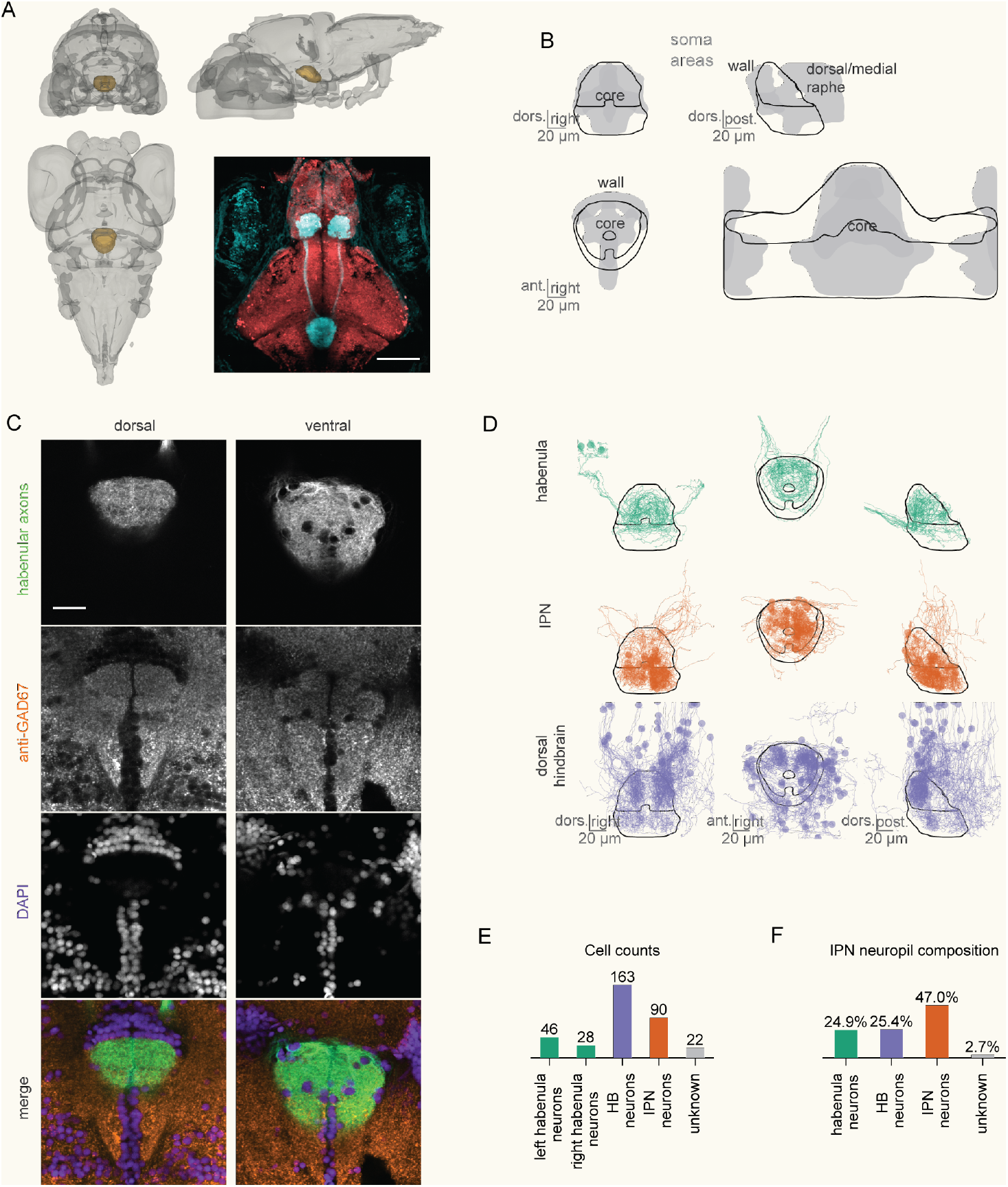
Cyto architecture of the IPN in larval zebrafish. A) Position of the IPN with in the larval zebra fish brain(MapZBrain Atlas); bottom right, an horizontal view of a triple transgenicfish Tg(16715:Gal4); Tg(UAS:GCaMP6s); Tg(elavl3:H2B-mCherry) in which GCaMP6s is expressed in the habenular neurons (cyan) and mCherry is expressed pan-neuronally (red). B) Distribution of somas around the IPN region. A central dense cluster of somas (core) separates the dIPN and the vIPN; a wall of somas delimitates anteriorly the dIPN. C) A horizontal confocal section of the dIPN (left) and vIPN (right) stained with anti-GAD67 (orange) and DAPI (purple). Hb axons (green) are labeled by Tg(16715:Gal4); Tg(UAS:GCaMP6s). D) Overview of the skeletons from EM reconstructions, separated by location of the cell somata (habenula, IPN core, dorsal hindbrain regions). E) Provenance of the cells reconstructed in this paper. F) composition of the IPN neuropil by cell provenance.

The habenula (Hb) neurons are considered as the major glutamatergic input into the entire IPN. While the existence of GABAergic neurons is reported in the IPN in larval zebrafish [43], the distribution of their axons and dendrites is not clear. We performed immunostaining against GAD67, and found that the distribution of the immunoreactive neuropils in the IPN overlap with the Hb axons (Figure 1C). Although the source of these GAD67-ir neuropils can also be other brain regions, this result shows that GABAergic neuropils densely cover the entire IPN.

### Morphological reconstructions of the habenulo-interpeduncular circuit

There is very limited evidence about the morphology of IPN core neurons (defined as neurons having the soma in the IPN). The only available data in larval zebrafish come from [44], which employed electroporation to label some IPN core neurons, of which the axon could not be observed. Therefore, we set out to investigate the cell morphologies of the region by manual skeletonization of cells from the volumetric EM dataset. We collected 349 complete or partial skeletons from neurons whose soma, dendrite or axon was within the IPN boundaries, as shown in Figure 1D (see Methods). First, we traced 90 of the 130 IPN core neurons (Figure 1E). Then, as the IPN circuit is highly participated by neurons in the dorsal hindbrain that elongate axons and dendrites in the region [39], we also reconstructed 163 neurons with either axon or dendrite in the IPN volume. Given the high participation of dorsal hindbrain cells to the IPN neuropil we decided to define any neuron with the dendrite in the IPN as an IPN neuron as they are functionally integrated in the IPN circuit. Finally, we reconstructed 76 complete or partial habenular neurons (31 complete cells and 45 just in the terminal part of the axon; 46 from the lHb, 28 from the rHb).

From our (non-dense) reconstructions, we could estimate that neurons with soma in the IPN core compose around half of the IPN neuropil (47%); habenular neurons contributed for around one quarter (24.3%); the rest was composed by the processes of anterior hindbrain neurons (25.4%) (Figure 1F).

### The habenulo-interpedunctular projection

The IPN has always been mainly defined by the massive projection it receives from the left and right Hb[refs]. Upon leaving the Hb, axons bundle in a very compact paired fiber tract, the fasciculus retroflexus (FR) (Figure 2A). By identifying the FR in the EM stack, we could precisely count the number of neurons composing the Hb-IPN projection in the 5 dpf fish, at two locations that were midway through the course of the tract. We could count 683-702 axons in the left FR and 555-581 axons in the right FR (Figure 2A).

**Figure 2:**
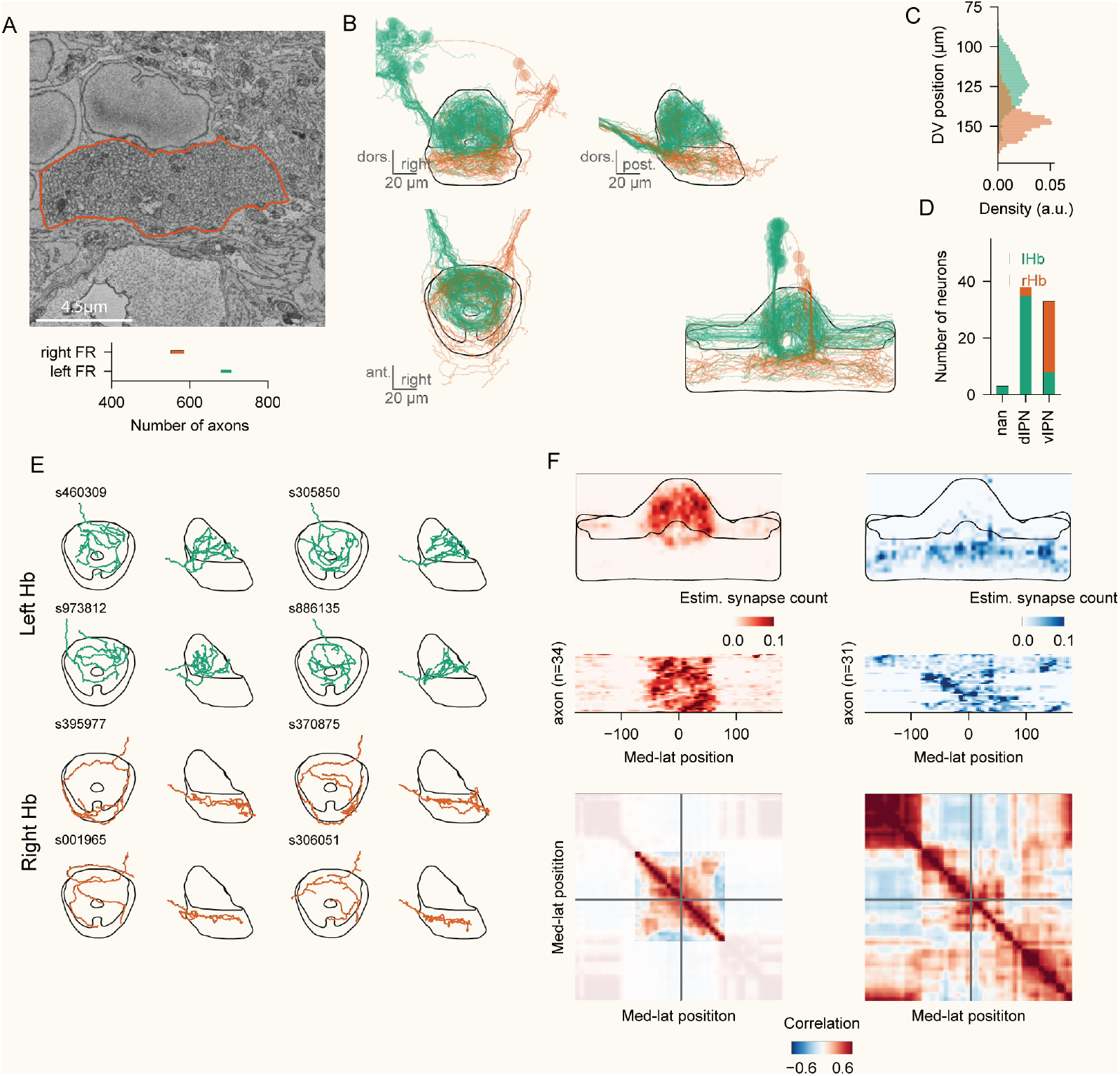
The habenulo-interpedunctular projection. A) Section of the right FR. Below, quantification of the number of axons in the left and right FR; the lines span the interval obtained by two axon counts at two different levels of the tract. B) Axons of the left (green) and the right (orange) Hb. C) Histogram showing the dorsoventral distribution of axons from the left and the right Hb. D) Counts of neurons from the left and the right Hb targeting the dIPN or the vIPN. E) Top and side view of individual axons from the left and the right Hb. F) Top: distribution of synapses around the IPN ellipse for dorsal (left) and ventral (right) axons. The middle panel represents for every axon the density of synapses along the ellipse. The bottom panel shows the cross-correlation of synapses distributions for every bin. This is a score that shows for every bin pair if neurons targeting the two bins are the same (high correlation) or different ones (low correlation).

We reconstructed 46 neurons from the lHb, and 28 from the rHb. For the incomplete reconstructions where the soma was missing, we inferred the provenance by looking at the FR from which the axon entered the IPN. As previously described [6], each axon targeted either the vIPN or the dIPN, but never both. As expected, we observed a strong tendency for left Hb neurons to target the dIPN (35/46 dIPN; 8/46 vIPN, 3/46 unassigned) and for right Hb neurons to target the vIPN (25/28 vIPN, 3/28 dIPN) (Figure 2B-D). The morphology of dIPN and vIPN projections was highly distinct, with dIPN Hb axons branching much more and covering the whole dIPN in a mesh-like arborization and vIPN axons spiraling in a concentric fashion over the vIPN with much sparser branching (Figure 2E).

### Distribution of synapses from the habenular projections

Do Hb axons provide unstructured input to the whole targeted IPN neuropil, or is there some organization principle in the Hb-IPN connectivity matrix? Previous investigations of the habenulo-interpeduncular projections were always limited by a resolution/coverage tradeoff, and could not provide a precise answer to this question over a large number of axons reconstructed in the same circuit. Leveraging the EM dataset, we manually annotated or estimated the localization of axonal terminals in the Hb projections in both the dorsal and the ventral IPN (see Methods).

In the dIPN (Figure 2E), the whole volume was homogeneously covered by synaptic terminals from Hb axons. Intriguingly, the distribution of contacts was not random: we could observe a striking pattern in the distribution of the axonal terminals from individual axons. Each neuron had a slightly biased concentration of boutons in some parts of the dIPN (Figure 2E). To better quantify this, we computed an “anatomical similarity matrix”, a measure of how similar two bins in the dIPN are, based on which neurons target preferentially the bins (Figure 2E). Remarkably, some neurons seemed to target preferentially both the left and the right side of the dIPN. This supports the notion that those apparently distant sides of the dIPN are close to each other in functional space as they represent the same heading direction, as proposed by [39].

In the vIPN, we observed a similar pattern (Figure 2E). Although the whole volume was covered by synapses from each Hb neuron, different axons seemed to target preferentially different areas in the vIPN. This was the case especially for the most caudal part of the region, that was often targeted preferentially by neurons that were not contacting the anterior part of the vIPN (unfortunately, the left portion of this caudal region was affected by a severe cut in the EM stack, which affected the tracing procedure).

### dIPN and vIPN neurons

While the organization of the Hb-IPN pathway is well established, little is known about the morphology and connectivity of neurons in the IPN that receive inputs from Hb axons. We therefore decided to investigate in further detail the morphology of cells with dendrites in either the dIPN or the vIPN, hereby referred to as dIPN neurons and vIPN neurons, regardless of the location of the soma (Figure 3A). We reconstructed the skeletons of 69 dIPN neurons, and 89 vIPN neurons. Overall, soma and dendrite were always on the same side of the brain for almost all cells (87/89 ipsilateral dendrites in vIPN neurons; 69/71 ipsilateral dendrites in dIPN neurons) (Figure 3A, top). The axon was often crossing the midline for about half the dIPN neurons (31/71 ipsilateral axons) and for one third of the vIPN neurons (57/89 ipsilateral dendrites) (Figure 3A, bottom).

**Figure 3:**
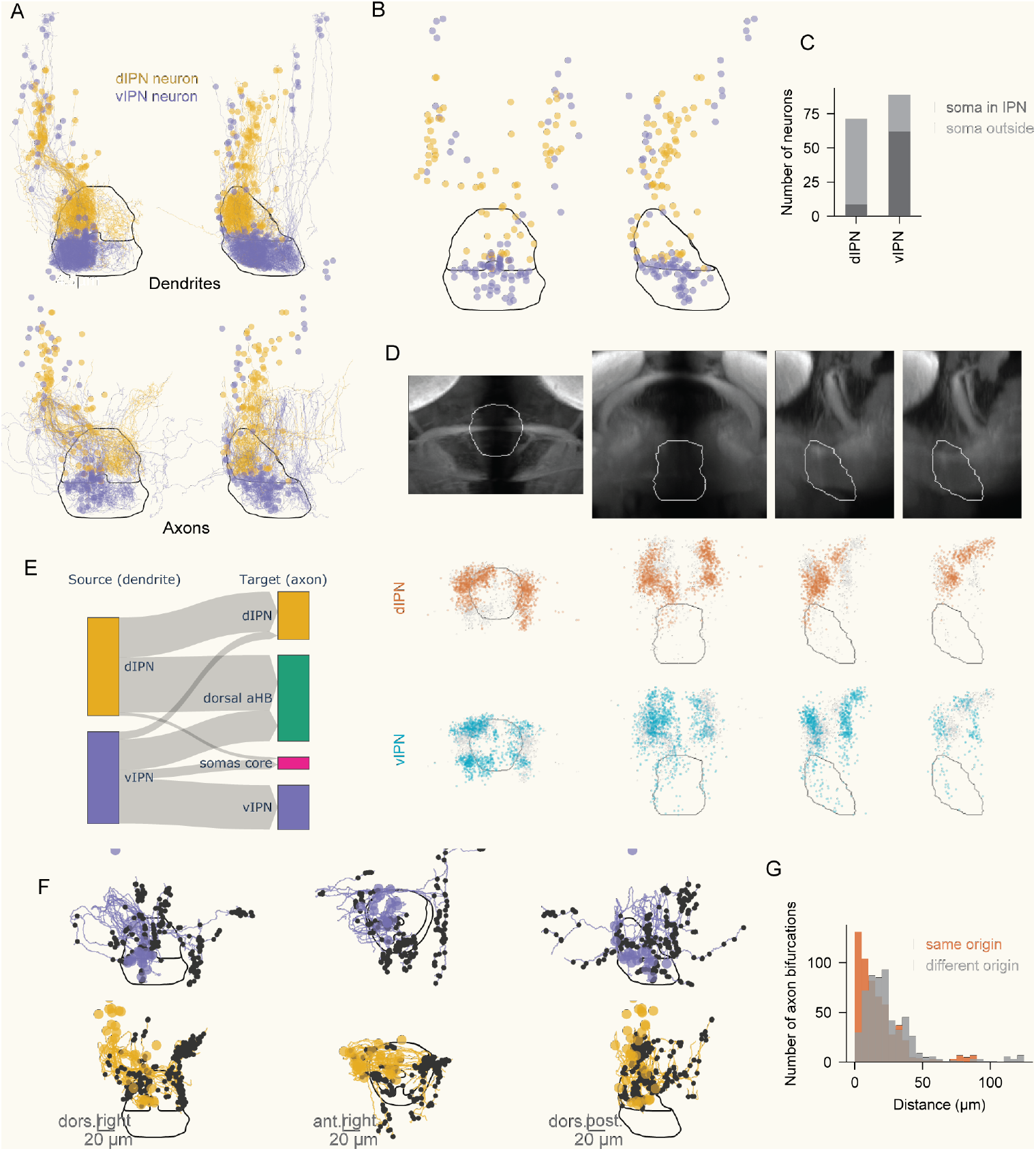
dIPN and vIPN neurons. A) Distribution of dendrites (above) and axons (below) for left-mirrored IPN neurons, defined as neurons with a dendrite in the IPN. Neurons are color-coded by the position of their dendrite (yellow: dIPN, blue: vIPN). B) Distribution of somas, color-coded by the dendrite position. C) Counts of neurons inside the IPN and outside the IPN with a dendrite in either the dIPN or the vIPN. D) Distribution of somas labelled by paGFP after photoactivation targeting the dIPN or the vIPN in the left hemisphere. The lateral view of the ipsi- (left) and contra-side (right) was shown separately. E) Sankey diagram for the projections of axons leaving the dIPN and vIPN. F) Distribution of axons of left-mirrored dIPN neurons (yellow) and vIPN neurons (blue) targeting the dorsal hindbrain. The black dots represent bifurcation points of the axons; as axons tend to branch out in their target region, such points are used as a proxy of synaptic density.

Previous observations from the lab [39] reported that a significant portion of the neuropil in the dIPN comprises dendrites of neurons whose somas lie outside the IPN, in the dorsal part of the first rhombomere (r1). To quantify this in further detail, we analyzed the soma location of all dIPN and vIPN neurons (Figure 3B-C). We confirmed that the large majority of dIPN neuron somas were in the r1 region above it (62/71 dIPN neurons), and almost none in the central core (2/71 dIPN neurons). On the other side, vIPN somas were mostly in the cell core (62/87 vIPN neurons), with some contributions from the dorsal r1 territory (27/87 vIPN neurons). Most dIPN and vIPN neurons with the soma outside the IPN shared the pseudounipolar morphology previously described for r1π neurons.

To better understand the distribution of vIPN and dIPN neuron somas, we performed photoactivation experiments using 6-8 dpf larvae ubiquitously expressing photoactivatable GFP (paGFP). Targeting the IPN neuropil regions, paGFP can be transported to the somas of the neurons that arborize in the IPN. We curated the location of the paGFP-labeled somas that are not in the IPN cell core or the Hb (Figure 3D); by targeting the whole or part of the dIPN and vIPN neuropils in the left hemisphere, 1359 cells from 40 animals and 1294 cells from 57 animals were labeled, respectively. vIPN labeled fewer cells in the contralateral hemisphere compared to dIPN. The most dorsal cells were detected right ventral to the cerebellum, and no cells more ventral to the IPN was detected. Although the distribution of the labeled cells was different between the dIPN and vIPN, all of the cells were labeled in the rostral hindbrain regions.

### Projections of IPN neurons

We then turned to investigate what regions received axons from IPN neurons. Here we refer to all IPN neurons with the axon targeting other regions of the IPN as IPN interneurons, and to all IPN neurons with the axon outside the IPN as IPN projection neurons. As is was not possible to carefully annotate synaptic terminals for all IPN skeletons, here and in the following analyses we use axon branching points distribution as a proxy for synaptic terminals, as those neurons tend to have high branching in the targeted region [39]. We excluded from the analysis neurons with very incomplete axons for which this classification was producing ambiguous results (see Methods).

Figure 3E shows the target regions for dIPN and vIPN neurons, identified as the region that contained the most axonal branching points for each neuron. Half (26/64) of the assigned dIPN neurons were dIPN interneurons, targeting different parts of the dIPN. Most of the other half (36/64) targeted the anterior hindbrain territory above the IPN. A similar pattern was observed in the vIPN: half of the assigned neurons (29/60 targeting the vIPN and 6/60 targeting the inner cell core) were vIPN interneurons, and one third (20/60) were vIPN projection neurons targeting other portions of the dorsal anterior hindbrain. Intriguingly, only a very small number of neurons projected from the vIPN to the dIPN (a small but morphologically consistent minority, see Figure 6); and none projected from the dIPN to the vIPN. This observation proves how the two subdivisions of the nucleus in larval zebrafish not only receive very different axonal innervation, but are also largely segregated in their inner connectivity.

Do projection vIPN and dIPN neurons target the same regions of the dorsal anterior hindbrain? When looking at the distribution of axon branching points outside the IPN for dIPN and vIPN neurons, they are largely non-overlapping (Figure 3F). vIPN neuron projections ended up mostly in a dorso-caudal region very close to the midline both contralaterally and ipsilaterally (presumably close to the raphe), or in a ventral region lateral to the vIPN (Figure 3F, top). dIPN neurons, on the other side, targeted mostly more lateral parts of the dorsal hindbrain, both contralaterally and ipsilaterally (Figure 3F, bottom). When looking at an axon branching point from an IPN projection neuron, its nearest neighbor was much more likely to come from a neuron of the same IPN subdivision (Figure 3G).

### Morphological clusters of dIPN neurons

We then proceeded to characterize thoroughly the morphological types that we observed in the IPN. First, we focused on the dIPN. By a combination of automatic criteria and manual annotation (see Methods), we converged to the definition of 7 morphological clusters for dIPN neurons (Figure 4A; Supplementary Video 1-7).

**Figure 4:**
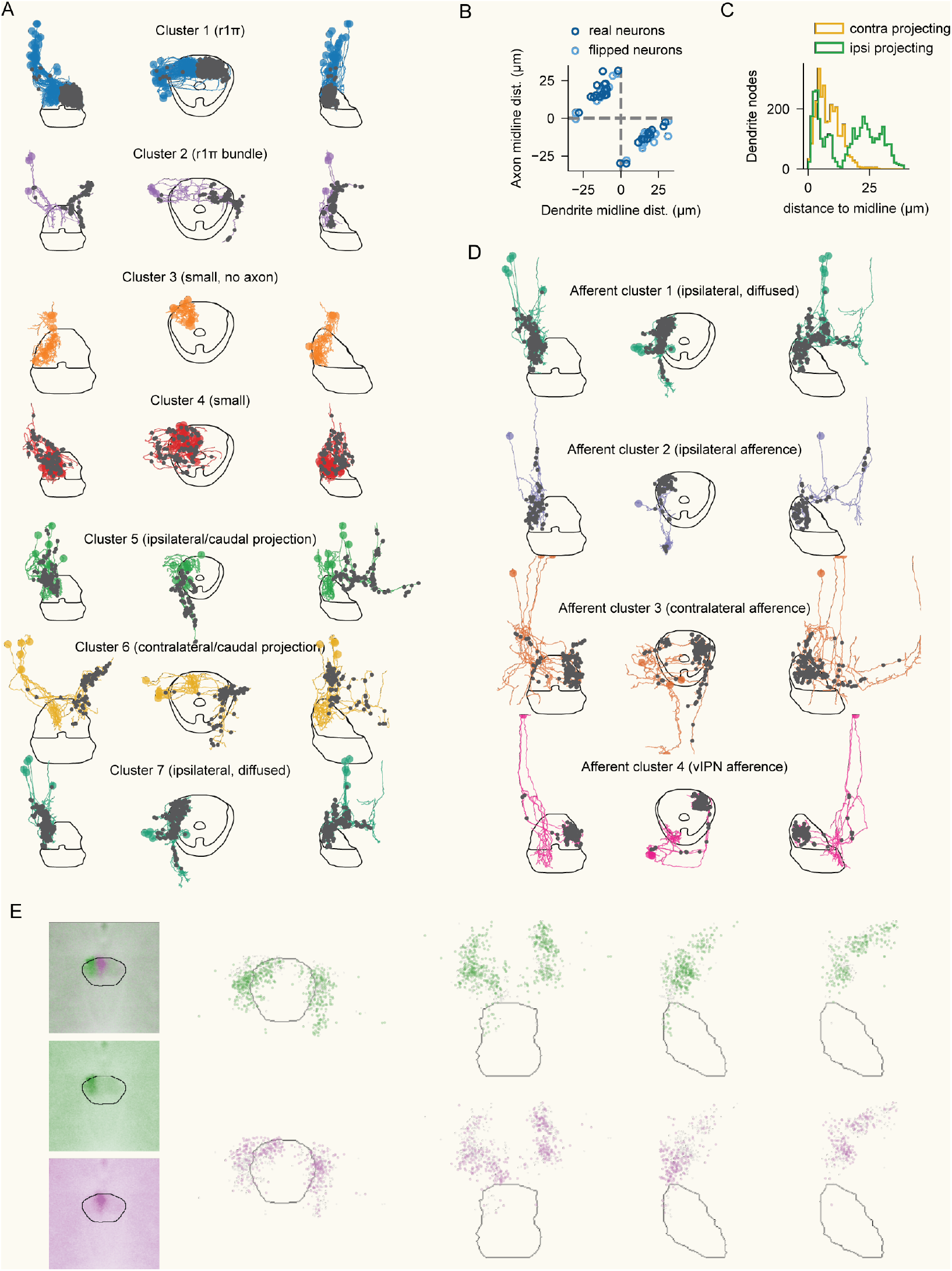
Morphological clusters of dIPN neurons. A) Each row shows different projections for all cells in a cluster; the dark dotsmark the axonal terminals ofthe cells. Seethe main text for the description ofthe clusters. B) Axon-dendritelocation relationship in r1π neurons: each dot represents the dendrite (x axis) axon (y axis) centroid for cells in the dataset (dark dots) and their flipped versions (light dots). Positions are tightly related with the previously described constant offset in dendrite and axon locations. C) Distribution of the dendrite nodes density around the midline for cells in clusters 6 and 7 shown in A. D) Morphological clusters in neurons projecting to the dIPN. Each row shows different projections for all cells in a cluster; the dark dots mark the axonal terminals of the cells. See the main text for the description of the clusters. E) Distribution of somas labelled by paGFP after photoactivation targeting the lateral or medial half of the dIPN in the left hemisphere. The lateral view of the ipsi- (left) and contra-side (right) was shown separately.

**Figure 5:**
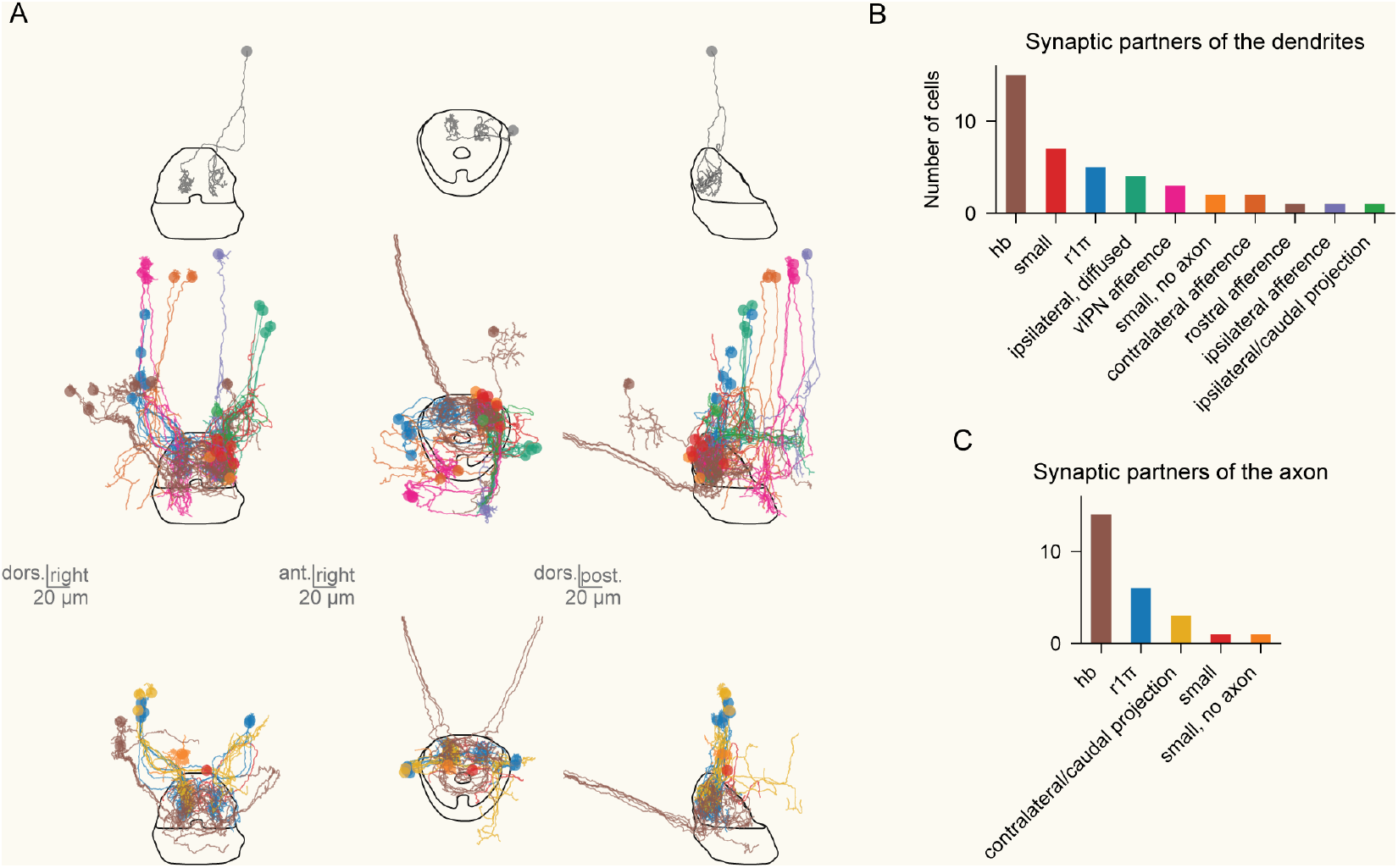
Connectivity of r1π neurons. A) Top: the focal neuron from which synaptic partners were reconstructed. Middle: partners of the dendrites, color-coded by cluster identity with the same color notation followed in Figure 4A. For clarity, only 5 habenular neurons are shown. Bottom: partners of the axon, color-coded by cluster identity. For clarity, only 5 habenular neurons are shown. B) histogram with the counts of different typologies for the partners of the dendrite. C) The same as B, for the partners of the axon.

The first dIPN cluster that we describe is the r1π type interneuron that was already observed in [39]. We could confirm the previously reported organization in axons and dendrites of r1π neurons, with a striking correlation between the location of the ipsilateral dendrite and the contralateral axon (Figure 4B). The second cluster consisted of cells very similar in morphology to r1π neurons, with an axon that extended both in the dIPN and dorsally in the hindbrain, following upward the bundle formed by descending processes of contralateral r1π neurons. The third cluster grouped cells that were characterized by a very compact dendritic tree very close to the soma, and the absence of an axon. The fourth cluster comprised similarly compact cells that also had a small axon that often terminated very close to the soma, or just dorsally above the IPN.

The last three clusters included dIPN projection neurons and were distinguished based on the target of the neuron’s axon. Cluster 5 included cells with a dorsocaudal ipsilateral projection, whole cluster 6 cells with a similar but contralateral projection. Interestingly, those cluster that were defined just on the basis of their axonal morphology were also clearly segregated in the distribution of their dendrites, with cells of cluster 5 having either very medial or very lateral dendrites, and cells of cluster 6 with dendrites in between the two peaks of cluster 5 dendrites density (Figure 4C). The last cluster included a group of cells where both the axon and dendrite were very extended rostro-caudally, and although they were clearly intercepting the dIPN, it was not the only region that they originated from or targeted.

Finally, we turned to afferent neurons that had their dendrite outside the dIPN and were targeting the dIPN with their axons (Figure 4D; Supplementary Videos 7-10). In our tracing effort we could not identify many neurons in this category, and the resulting clusters are not as clear-cut and delineated as the clusters for dIPN neurons. The first cluster corresponded to cluster 7 of dIPN neurons. The second cluster included neurons with a small and poorly branched dendrite in the dorsal hindbrain, and an ipsilateral projection to the dorsal IPN. The third cluster was composed of neurons with an ipsilateral dendrite in the dorsal hindbrain, and a contralateral axon in the dIPN. The last cluster included the neurons that projected to the dIPN from the caudal vIPN.

To further characterize the distribution of the cells arborizing in the dIPN, we performed paGFP photoactivation targeting the lateral and the medial half of the dIPN neuropils in the left hemisphere (Figure 4E). The regions in which the cells were labeled by targeting the lateral (519 cells from 15 fish) and medial (338 cells from 12 fish) dIPN were similar, while the density was different. The lateral dIPN labeled more cells in the ipsilateral hemisphere, whereas the medial dIPN labeled more on the contralateral side. On the ipsilateral side, the medial dIPN seemed to label slightly more ventral cells than the lateral IPN. These results suggest that the dIPN has a certain anatomical structure along the medio-lateral axis, although it is yet unclear whether the dIPN can be divided into distinct stripes or it is rather a gradient.

### Connectivity of r1π neurons

r1π neurons seem to be the most common morphology in the dIPN. Moreover, it has been suggested that r1π neurons could integrate a ring attractor network that accumulates directional movement to compute heading direction representations Therefore, we decided to characterize in detail the distribution of synaptic partners of a chosen r1π neuron.

We identified the synaptic partners of the axons and dendrites of the chosen neuron from the EM dataset (Figure 5), and clustered them based on the connectivity and the morphology as described in Figure 4. The majority of the partners were Hb axons for both the axons and dendrites of the chosen neuron via axo-axonic and axodendritic synapses, respectively. The second largest cluster was the r1π neurons, which again confirmed our previous observation [39] supporting the reciprocal connectivity between themselves. Another abundant cell type is the cluster 3 and 4 (small cells), often located immediately rostral to the dIPN neuropil region and formed dendro-dendritic synapses with the chosen neuron. One noticeable presynaptic partner is the afferent cluster 4 (caudal vIPN to the contralateral dIPN) located on the contralateral side, which is the small population that directly connects vIPN and dIPN (See Figure 3E). One prominent cell type of the postsynaptic partners was the cluster 6 (contralateral caudal projection) located on the contralateral side of the chosen neuron. These results demonstrate that various cell types are potentially involved in the ring attractor network via canonical and non-canonical synapses, whereas their roles in the circuit remain to be elucidated.

### A glomerular organization for the vIPN

We then turned to the investigation of the vIPN. We started by noticing that the vIPN cells were categorizable using their dendritic tree density, and we divide them in cells with dense dendritic trees (54/87 neurons), and cells with sparse dendritic trees (33/87 neurons) (Figure 6A). Cells with sparse dendritic trees often had an early-terminating and poorly branching axon (Figure 6B, left). For this reason we suspected that the sparsely branching neurons corresponded to cells that were still in maturation; as we could not find clear clusters in their morphologies, we focused our analysis on cells with dense dendrites (Figure 6B, right).

When looking at the distribution of dendrites from the whole population of dense vIPN neurons, we could observe six clear clusters of high vIPN dendrites densities around the ellipsoid of the vIPN (Figure 6C), three in the left vIPN and three in the right vIPN. When looking back at histological data, we realized that these clusters can also be appreciated in the GAD67 immunoreactive neuropils (Figure 1C). The distribution of the Hb axons and GAD-ir neuropils in the IPN largely overlap, while gaps in the GAD-ir neuropils could be observed in the midline of the dIPN and between these six clusters in the vIPN.

When looking at the distribution of dendrites from individual neurons, it was clear how each neuron had a dendrite that was branching extensively and exclusively into one of those densities (Figure 6B). For this reason, we decided to name those density glomeruli: highly packed densities of neurites approx. 70 microns in diameter, sharing common boundaries. Intriguingly, the glomeruli form an evenly spaced structure in the vIPN, transversed by the crossings of left Hb axons (Figure 2E). The rostral glomeruli (left/right glomerulus 1) were the densest and smallest, caudal glomeruli (left/right glomerulus 3) the least dense and broadest. Intermediate glomeruli (left/right glomerulus 2) had density and measures in between (Figure 6B). We refer to neurons with a dendrite in any of the glomeruli as glomerular neurons of the vIPN.

The population of glomerular neurons included both projection neurons and vIPN interneurons. vIPN interneurons (Figure 6D), were all targeting each one a specific glomerulus with their axon; moreover, the (source) dendrite glomerulus was highly predictive of the target glomerulus. Glomerulus 1 interneurons were all targeting contralateral glomerulus 2 (Figure 6D, top). Glomerulus 2 interneurons were mostly targeting contralateral glomerulus 1; some of the neurons targeting controlateral glomerulus 1 were also branching in the ipsilateral glomerulus 1. Other glomerulus 2 interneurons were targeting the ipsilateral or the contralateral glomerulus 3 (Figure 6D, middle). Finally, glomerulus 3 interneurons were all targeting the contralateral glomerulus 3 (Figure 6D, bottom).

vIPN glomerular projection neurons also had very distinct projection patterns outside the vIPN depending on their source glomerulus (Figure 6D). Glomerulus 1 neurons were extending axons dorsally, either close to the midline (both ipsi- and contralaterally), or laterally, far from the midline, in the contralateral side (Figure 6D, top). Glomerulus 2 neurons were all projecting dorsally and ipsilaterally (Figure 6D, middle). Glomerulus 3 neurons all projected to the dIPN (Figure 6D, bottom).

To further understand the source of the projections in the vIPN, we performed paGFP photoactivation targeting each vIPN glomerulus in the left hemisphere (Figure 6E). The cells were labeled in distinct regions with partial overlap by targeting the medial (407 cells from 25 fish), lateral (379 cells from 18 fish), and caudal (508 cells from 14 fish) glomerulus. The medial glomerulus labeled the most rostral regions of the hindbrain bilaterally, and the caudal glomerulus labeled more caudal and also dorsal population bilaterally. The lateral glomerulus labeled both the rostral and caudal regions overlapping with the other two glomeruli, but mostly in the ipsi-lateral hemisphere. These results confirm the observation of the projection patterns of vIPN interneurons (Figure 6D), and also suggest that each glomerulus has its unique source of projections from the rostral hindbrain regions.

**Figure 6:**
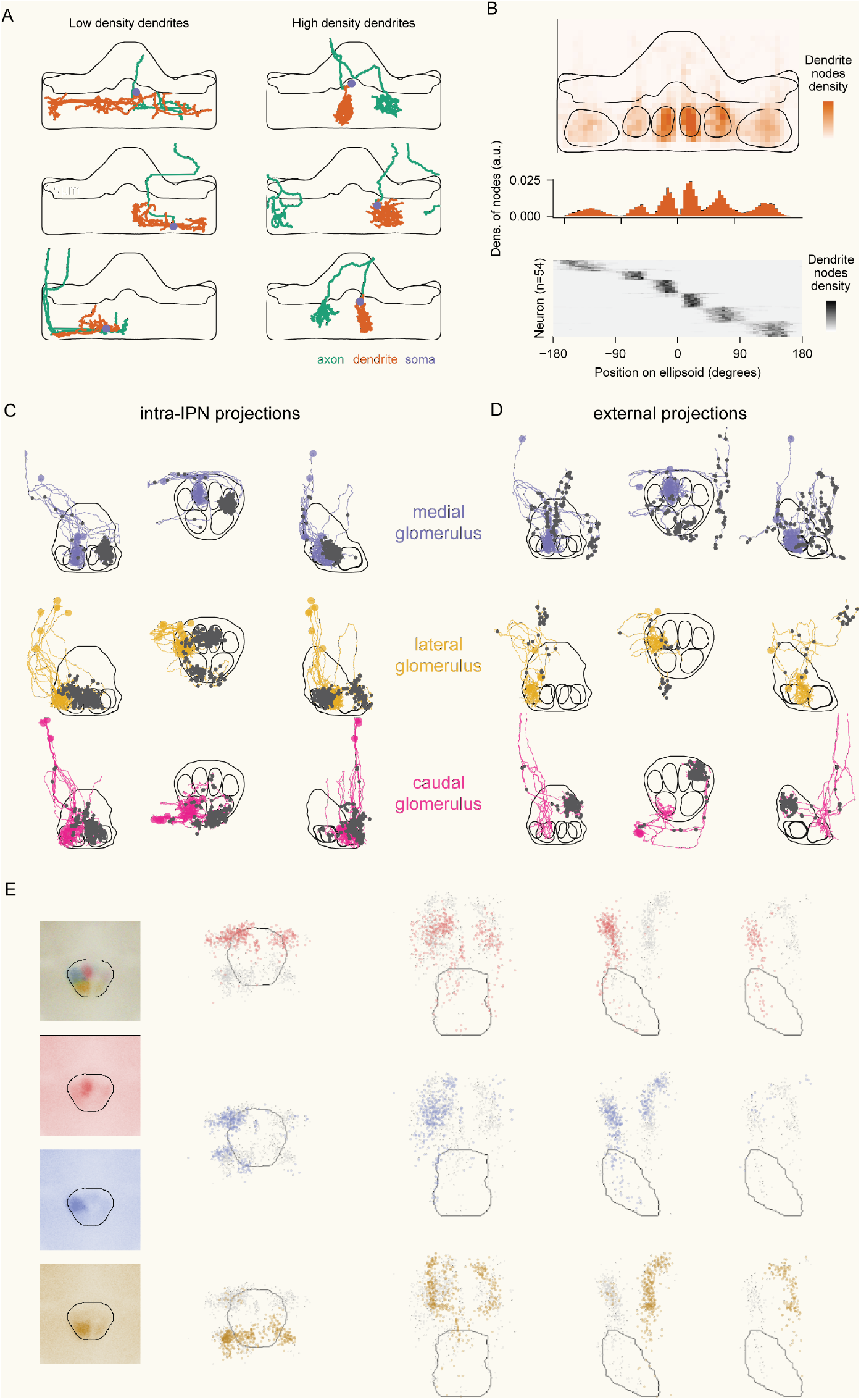
A glomerular organization for the vIPN. A) Examples of individual neurons with low (left) and high (right) density dendrites. B) Top: distribution of dendrite notes for all neurons with high density in the vIPN visualized over the IPN ellipse projection, with the glomeruli outlined. Middle: histogram with node density along the ellipse length. Below: dendrite node density of individual neurons, sorted by their center of mass. There is a clear discrete separation between different glomeruli. C) Glomerular neurons (left-mirrored) separated by their dendrite glomerulus and by their targeting: left, neurons projecting to other glomeruli in the vIPN; right, neurons targeting other regions. Black dots represent axon branching points. D) Distribution of somas labelled by paGFP after photoactivation targeting the medial, lateral, orcaudal glomerulus of the vIPN in the left hemisphere. The lateral view of the ipsi- (left) and contra-side (right) was shown separately.

### Hb axons calcium activity is shaped by vIPN neurons morphology

Having elucidated the inner structure of the IPN neuropil beyond the known projection patterns of Hb neurons, we turned to the analysis of a calcium imaging dataset acquired from the axons of the Hb neurons expressing GCaMP6s in the IPN. The spontaneous activity in the dark was recorded with two-photon microscopy in a plane, and the analysis was done using the periods in which no obvious tail movement was observed. To characterize the pattern of the population activity, we computed a correlation matrix by splitting the Hb axons into 30 bins along the medio-lateral axis and the ellipsoid for the dIPN and vIPN, respectively. In the dIPN, the bins that are anatomically close to each other show high correlation, and the bins at the left and the right ends also tend to show high correlation to the less extent (Figure 7A, E). In the vIPN, the ellipsoid can be roughly divided into six anatomical clusters that are functionally distinct from each other (Figure 7B, F); bins that are located within a certain anatomical position are highly correlated, whereas two adjacent bins can also be anti-correlated if they belong to the different clusters. These activity patterns differ from what would be expected from the anatomical localization of each Hb axon in the IPN (Figure 2), where no strong bias in the distribution was noticed either in dIPN or vIPN.

**Figure 7:**
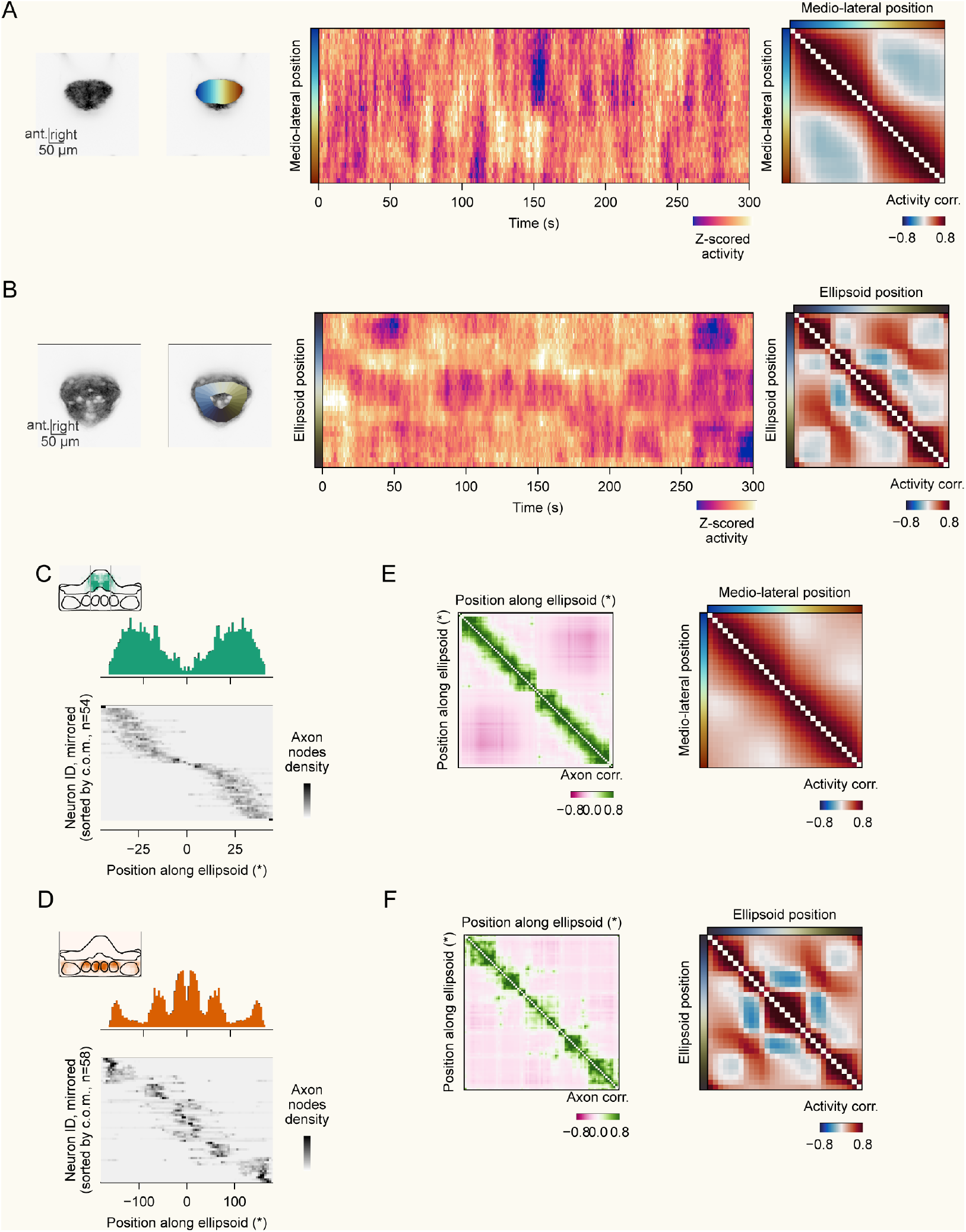
Spontaneous calcium activity in Hb axons. A) An example anatomy and bins of the Hb axons expressing GCaMP6s in the dIPN, a five minutes period of the spontaneous activity recorded in the dark, and the average correlation matrix of the bins from the animal. B) Same as A for the vIPN. C) Distribution of the axons of IPN neurons projecting inside the dIPN. Top: cumulative histogram; bottom: neuron-wise histogram. The neurons were mirrored. D) Same as D, for the vIPN. E) Left: Bin-wise correlation of dIPN neurons axons targeting different bins of the dIPN. The binning is done over the ellipse projection described in Figure 1S and it closely matches the binning performed over the imaging data. Right: correlation matrix of activity in different medio-lateral sectors of the dIPN, with the binning shown in Figure 7A. The matrix is the average over n=43 fish. F) Same as E, for the vIPN.

To quantify the degree to which this activity could be due to the organization of axons from IPN interneurons, we turned our attention back to the distribution of the axons from dIPN and vIPN interneurons. In the dIPN, the interneuron axons tiled homogeneously the whole medio-lateral extension of the nucleus, with no clear clusters apart from the left/right pattern resulting from the axons of individual cells never crossing the midline (Figure 7C). In the vIPN, however, the glomerular structure that we described above produced a clear segregation of axons in individual glomeruli, with the only notable exception of left and right glomeruli sometimes being both targeted by the same axon (Figure 7D).

To better quantify those patterns and compare them with the functional correlations we observed, we computed a “axon similarity matrix” that, for every pair of bins along the IPN ellipse, was providing a metric of how much the pair was targeted by the same or different axons. When put side by side with the functional correlation matrices, the two metrics showed very consistent patterns: a continuous distribution of high correlations with the neighbor bins in the dIPN, with a small discontinuity close to the midline (Figure 7E); and segregated blocks of high correlation for individual glomeruli for the vIPN, with the exception of the two first glomeruli that exhibited high correlation in the functional imaging dataset and shared axonal input in the input similarity matrix (Figure 7F). Note that this analysis does not include any information about the distribution of the dendrites associated with the axons. We believe that certain parts of the imaging correlation structure (such as the increased correlation of bins at the two left and right extremes of the dIPN) would require using dendrite distribution information to be explained (namely, the similar placing along the midline for r1π neurons targeting the left and right extremes of the dIPN). More careful analyses and simulation will produce further insights into how structure and function interact in the IPN.

## Discussion

In this work, we provide the first thorough characterization of the habenulo-interpeduncular circuit in larval zebrafish using a combination of histology, EM volumetric reconstruction, targeted cell labelling, and two photon imaging.

First, we confirmed and expanded upon the previous characterization of the habenulo-interpeduncular projection [5, 6]. We observed that, beyond the known left-right asymmetry of the habenular projections to dIPN and vIPN [6, 45], a second level of organization could be found at the level of synaptic terminals. We observed that, although each axon made synaptic contacts across the IPN territory, there were consistent biases in the tendency to make contacts in different IPN locations. Intriguingly, those biases overlapped with the functional and anatomical connectivity of downstream IPN circuits. In the dIPN, habenular synapses from each neuron were denser in specific locations of the dIPN medio-lateral axis; remarkably, axons with denser synapses on the lateral-most portion of the dIPN had also a contralateral density in a similarly lateral location, consistently with the topographical organization previously described for r1π neurons, a subtype of dIPN interneurons [39]. Similarly, in the vIPN, the densities of habenular axon terminals reflected to some degree the glomerular organization of the neuropils that consist of local IPN neurons and other hindbrain neurons.

Next, we refined and extended previous observations on IPN cell types [6, 7, 9, 39]. We confirmed the topographical arrangement of r1π neurons, and we show how the IPN internal circuitry extensively involves other neurons whose soma is outside the IPN, especially in the dIPN circuit. Many neurons located in more dorsal regions of the hindbrain extend both their dendrite and axons into the IPN, and should therefore be considered as IPN interneurons. Similarly, several projection neurons targeting downstream regions have their soma located outside the IPN. Our observations complement existing reports on the functional diversity of cells whose somas are intermingled in the highly soma-dense region of the larval zebrafish hindbrain [46–49]. Moreover, they raise the intriguing possibility that the existing data coming from bulk anterograde and retrograde labeling of tegmental and interpeduncular cells in the mammalian IPN [4, 12] could result from similar, yet underscribed cell types contributing both dendrite and axons to the IPN.

For the first time, we could describe a strict segregation of the dIPN and the vIPN circuit in larval zebrafish. Although it was known that the two IPN subregions receive strictly segregated projections from different subdivisions of the habenula, the degree of segregation in the downstream circuit, consisting of the dIPN and the vIPN was not known. Our EM reconstructions showed that pairs of IPN neurons with dendrites in the dIPN and the vIPN respectively have almost no reciprocal connections, with the exception of a cell type projecting from the caudal vIPN to the contralateral dIPN. Furthermore, vIPN and dIPN projecting neurons targeted non-overlapping brain regions in the dorsal hindbrain. This segregation suggests that the dIPN and the vIPN should be considered as separate functional units, each receiving input from a different habenula subnucleus, and processing information with independent circuitry, potentially mediating different functions with similar circuit motifs.

We also described a prominent glomerular organization of the vIPN. There, we observed six strictly compartmentalized glomeruli, approximately 20-30 μm in diameter, organized in a circle around the vIPN. Moreover, we show how, in each glomerulus, dendrites or axons from different IPN cell types branched extensively within some shared boundary, likely as the result of local morphogenetic cues and cell-cell surface interactions. Previous observations have hinted at the existence of this cell morphology; moreover, evidence from EM on the mammalian IPN has suggested the existence of similarly organized glomeruli, defined as dendritic trees of IPN cells ensheathing a large number of habenular axons [50]. With our volumetric reconstructions, we observe clear connection patterns across glomeruli: for example, a strong reciprocal connectivity between the first, anterior glomerulus and the contralateral, lateral glomerulus. Considering the GABAergic nature of most IPN neurons[43], those reciprocally inhibiting glomeruli could implement reciprocal inhibition of inhibition, a circuit motif that has already been described for processing competing stimuli [51, 52], and for action selection [53, 54].

Importantly, vIPN glomeruli are extensively traversed by habenular axons looping around the vIPN region, suggesting a computational module where information carried by the habenular axons is broadcast to different glomeruli, which could integrate it and mutually suppress with each other thereby resulting in different behavioral outcomes. This overarching organization is coherent with the decision-making function that is being highlighted for the IPN from many different perspectives and paradigms [17, 30, 41, 55]. Under this hypothesis, the slightly biased distribution of habenular axonal terminals in different glomeruli could reflect plasticity mechanisms in action to potentiate or suppress the synapses that target each glomerulus, similarly to what happens in the parallel fibers traversing the dendritic trees of Purkinje cells in the cerebellum [56] or to ring neurons targeting the ellipsoid body in *Drosophila* [57, 58]. Such potentiation has already been reported in mammals [59], and could act in synergy with short term plasticity [60] post-synaptic plasticity mechanisms that have been recently suggested to support decision-making computation in the IPN [31]. Indeed, by imaging calcium in the axons of habenular neurons we could observe a striking compartmentalization in their spontaneous activity. Our data show how the activity of anatomically widespread habenular projections to the IPN could actually be orchestrated by the local circuitry, with a clear correspondence between ongoing calcium dynamics of the habenular axons and the anatomical architecture of the IPN neuropils. This observation is consistent with the known effect of GABA release from IPN neurons on GABA_B_ receptors on habenular axons, reported in both fish [43] and mammals [59–62]. Such a conservation hints at a fundamental role of this presynaptic modulation in the function of the habenulo-interpeduncular pathway. The anatomical organization of axo-axonal modulation we describe could afford the integration of habenular inputs, afferences from other regions, and ongoing activity in the IPN already at the level of presynaptic release.

Overall, our work sheds light on the internal organization of a poorly understood brain region that is increasingly being implicated in a number of cognitive functions. We confirm and refine previous observations on the organization of the dIPN, and describe new circuit motifs that could underlie information processing in the vIPN. Moreover, we provide the first characterization of how previously described axo-axonal interactions in the IPN orchestrate the local calcium dynamics of habenular axons. The highly conserved nature of the habenulo-interpeduncular pathway suggests that these circuit motifs could also exist in the mammalian IPN. Future experiments will undoubtedly provide us with a more thorough understanding of the functional meaning of such a remarkable organization of the IPN.

## Acknowledgements

We thank all the members of the Portugues lab for their input. This research was funded by the German Research Foundation (DFG) under Germany’s Excellence Strategy within the framework of the Munich Cluster for Systems Neurology (EXC 2145 SyNergy, identifier 390857198) and through the “Enhanced resolution microscopy” project DFG – Projektnummer 518284373, and by the Volkswagen Stiftung via a Life? grant.

## Materials and Methods

### Zebrafish husbandry

All zebrafish handling procedures complied with protocols sanctioned by the Technische Universität München and the Regierung von Oberbayern (animal protocol number 55-2-1-54-2532-101-12 and 55.2-2532.Vet_02-24-5). Adult zebrafish (Danio rerio) of the Tupfel long fin strain were maintained at a temperature of 27.5–28°C under a 14-hour light and 10-hour dark cycle. They were housed in a fully recirculating water system with carbon, bio-, and UV filtration, and 12% of the water was exchanged daily. The water pH was maintained between 7.0 and 7.5 with a 20 g/l buffer, and the conductivity was kept at 750–800 µS using 100 g/l. Fish were kept in 3.5-liter tanks in groups of 10 to 17. Adults were fed twice daily with Gemma Micro 300 (Skretting) and live Artemia salina, while larvae received Sera Micron Nature (Sera) and ST-1 (Aquaschwarz) three times daily.

Experiments were conducted on 6 to 9 days dpf larvae of undetermined sex, except from the EM volume acquisition for which a 5 dpf larvae was used. Prior to the experiments, breeding was set up with one male and one female or three males and three females overnight in a Tecniplast Sloping Breeding Tank. The next morning, eggs were collected, rinsed with water from the facility system, and kept in groups of 20 to 40 in 90-cm petri dishes containing 0.3× Danieau solution (17.4 mM NaCl, 0.21 mM KCl, 0.12 mM MgSO_4_, 0.18 mM Ca(NO_3_)_2_, 1.5 mM HEPES; reagents from Sigma-Aldrich) until hatching. Post-hatching, larvae were maintained in water from the fish facility in an incubator set to 28.5°C and a 14-hour light and 10-hour dark cycle, with the solution being changed daily. At 4 or 5 dpf, larvae were lightly anesthetized with tricaine mesylate (Sigma-Aldrich) and screened for fluorescence under an epifluorescence microscope.

### Immunostaining

Immunostaining and imaging were performed on whole mount brains. Tg(16715:Gal4) [41]; Tg(UAS:GCaMP6s)mpn101 [63] double transgenic larvae were fixed at 6 dpf with 4% w/v Paraformaldehyde (Sigma) in 1x PBS (pH 7.4; 137 mM NaCl, 2.7 mM KCl, 8 mM Na_2_HPO_4_, 1.5 mM KH_2_PO_4_) at 4°C overnight. The brains were dissected from the fixed animals and incubated with the blocking solution (0.3 % v/v Triton X-100, 4 % w/v bovine serum albumin in 1x PBS) at room temperature for 3 hours. The brains were incubated with a rabbit anti-GAD67 antibody (PA5-21397, Invitrogen) in the blocking solution (1:300) at 4°C for 3 days, and then with an anti-rabbit IgG antibody conjugated with Alexa Fluor 568 (ab175470, Abcam) in the blocking solution (1:500) at 4 °C for 3 days. The brains were mounted in 0.7 % w/v low melting point agarose (Invitrogen) in 1x PBS, and imaged with a confocal microscope FV1000 (Olympus) using a 25x water-immersion objective (NA 1.1, Nikon).

### Photoactivation of photoactivatable GFP

At 6-8 dpf, Tg(Cau.Tuba1:c3paGFP)a7437 [64]; Tg(elavl3:LY-TagRFP)mpn404 [65] or Tg(elavl3:H2B-mCherry) [39] double transgenic larvae were mounted in 2% agarose, and placed under a custom-built two-photon microscope. The region for photoactivation was selected by imaging the pan-neuronal TagRFP or mCherry signal excited by a 950 nm laser. Photoactivation was performed at 800 nm, by six repetitions of 10 second continuous scanning with a 20 second break between each repetition. The laser power was 14 mW measured at the objective back aperture. 4-5 hours after photoactivation, the whole brain was imaged under a confocal microscope FV1000 (Olympus) using a 20x water-immersion objective (NA 1.0, Olympus). The location of the somas labeled by paGFP was annotated manually, and the images were registered to a reference brain using CMTK [66].

### Two-photon imaging and analysis

Functional imaging of the habenular axons in the IPN was done using Tg(16715:Gal4); Tg(UAS:GCaMP6s) animals at 6-8 dpf with a custom-built two-photon microscope controlled by a Python package Brunoise [67]. The animals were embedded in 2% w/v low melting point agarose (Invitrogen) in the water from the fish facility. The images were acquired at 3 Hz for 30 minutes or longer for each plane using 905 nm laser, and the tail movements were tracked by a Python package Stytra [68] under the illumination with 830 nm LED. Thirty seconds before and after the frames at which tail movements were observed were removed from the recorded time series, and the resulting periods longer than three consecutive minutes were used for the analysis. The IPN was split into 30 bins along the medio-lateral axis for the dIPN or along the ellipsoid for the vIPN. A correlation matrix was computed for each period, and averaged over the periods for each animal.

### EM reconstructions

EM reconstructions were performed on an open-access Serial Blockface EM dataset obtained from a 5-dpf larval Tg(elavl3:GCaMP5G)a4598 zebrafish sampled at a resolution of 14 × 14 × 25 nm (described in [42]). The initial identification of the IPN site within the EM stack wassuggested by the distinct arrangement of neuropil and cellular bodies in the ventral area of rhombomere 1. Subsequently, this identification was verified by following axons traceable to the habenulae via a fiber tract clearly recognizable as the fasciculus retroflexus due to its trajectory.

Skeletonization was conducted manually by annotators at ariadne.ai AG starting from seed points chosen by the authors. Upon completion of a cell, an expert annotator conducted a quality review. Difficult areas were then assessed by the expert, who would return them to the annotator team for further tracing if required. This process was repeated until all cells were completely traced. The annotators, including experts from Ariadne or the authors, independently labeled the skeletons to identify dendrites and axons based on morphological characteristics such as process thickness and the presence of presynaptic boutons, achieving consistent results. Half of neurons in the dataset were reconstructed by annotators by merging together neurite segments obtained by the automatic flood-filling algorithm described in [42]. Those neurons underwent the same quality control than the manually skeletonized neurons. From the aggregated mesh, the skeleton was then extracted using the *skeletonize*.*by_wavefront()* function from the skeletor library (version 1.2.3) and further refined using custom Python scripts. All subsequent analyses and quantifications of the reconstructions were executed using Python.

The dataset was obtained in subsequent rounds of tracing starting from seeds chosen with different criteria, namely: random seeds placed in neurons from the IPN core (79/349 neurons); random seeds placed in processes in dIPN and vIPN (37/349 neurons); seeds placed in neurons identified as potential candidates to be r1π neurons, as described in [39] (17/349 neurons); random seeds placed at one plane of the left and right FR (41/349 skeletons); neurons identified as the synaptic partners of r1π neurons (175/349 skeletons).

### IPN atlas definition

High-resolution image stacks of the IPN were collected from 7 dpf zebrafish larvae across various transgenic strains (Tg(16715:Gal4) [41]; Tg(UAS:NTR-mCherry)c264 [69], Tg(elavl3:H2B-GCaMP6s)jf5 [70], and TgBAC(gad1b:GFP)nns25 [71]). Imaging was performed using a 20× water immersion lens (NA = 1.0) with voxel dimensions of 0.5 μm × 0.5 μm × 0.5 μm using an LSM 880 microscope from Carl Zeiss. These stacks were aligned using the dipy library [72], focusing on the channel displaying the Tg(16715:Gal4) signal.

Subregions of the IPN were manually delineated within the volumes using the napari imaging tool. Then, an affine transformation from the EM data coordinates to the IPN space coordinates was obtained manually, matching the boundaries of the IPN and the glomeruli in the two datasets.

### Flattened IPN projection

To obtain the flattened projections of IPN neurons, we manually defined an elliptical cylinder centered around the IPN center, and intercepting all six glomeruli (see Figure S1). The projection process was implemented by designing a dense 2D square grid on the ellipse, with an (approximately) isometric size of 0.5×0.5 um when locally flattened; each point to be projected was then transformed into the nearest grid point of the ellipse. The transformed coordinates are shown using a new arbitrary x axis going from -180 to +180 mapping the whole perimeter of the ellipse.

### Synapses / branching points definition

In Hb axons, where the axonal terminals were very clearly distinguishable as enlargements in the diameter of the axonal process with synaptic vesicles puncta inside, we used two methods to obtain the coordinates of axonal boutons. For the vIPN axons, which were not branching extensively, we manually labeled all axonal boutons occurrences. For dIPN axons, we used the diameter of the mesh as a proxy for the presence of synaptic boutons; this proxy was validated by manual inspection of some of the sites identified as boutons in the EM dataset.

For the other neurons in the dataset, we could not annotate manually all the synaptic terminals, nor infer them automatically. Therefore, we relied on the observation that axons were mostly branching at the target region where most axonal terminals could be observed. We defined axonal branching points as the nodes of the skeleton with more than 2 edges; and located in the axonal part of the skeleton. Those are the points used in the visualizations of IPN cells axons. Similarly we defined dendrite branching points.

### Visualization

The skeleton was visualized using custom scripts and notebooks in Python, using matplotlib. The whole brain visualization in Figure 1 was created using brainrender [73] and the mapZbrain atlas [74]. The videos of the reconstructed neurons were created using Blender (Blender Foundation) with the polygon meshes from the EM dataset and the whole brain mesh from the mapZbrain atlas.

### Criteria for the classification of neurons

Neurons in the dataset were classified using a combination of manual annotation and automatic criteria.

#### Soma classification and mirroring

Somas were classified as being in the IPN if their coordinates were located in the whole IPN mesh. Cells were classified as left or right depending on their mediolateral position in the IPN reference coordinates. In mirrored panels, right cells were just flipped along this axis.

#### Habenular neurons

All skeletons that consisted of axons descending from the FR and arborizing in the IPN were considered to be Hb neurons. Of the 76 Hb neurons described in this way, 31 were fully reconstructed up to the soma. No consistent differences were found in the axonal morphology of neurons with and without a fully reconstructed soma.

#### IPN and hindbrain neurons

Neuron processes (axons and dendrites) were defined as belonging to a given region or subregion of the IPN if the majority of their nodes were contained within that region. dIPN neurons (neurons with the dendrite in the dIPN) were further classified by manual annotation into the clusters presented in Figure 4 looking at their morphologies. vIPN neurons were first classified into sparse and dense dendrites groups based on a threshold as their dendrite density (computed as median distance between dendrites nodes); then, dense dendrite cells were divided based on the glomerulus that contained their dendrite.

### Anatomical similarity correlations

For the anatomical similarity matrices of Figure 2F and Figure 6E-F, we first created for every neuron an histogram of node distribution along the IPN projection ellipse. Then, we put together the (normalized) histograms for all neurons in a (B, N) matrix (with N number of neurons and B number of ellipse bins). From this matrix we computed the (B, B) correlation matrix, where each entry describes the correlations between the N-dimensional vectors of neuron densities in each of the two bins.

## Data and code availability

All the data described in the paper, the analysis notebooks and the figure generation notebooks will be made available upon publication.

## Supplementary videos and figures

**Supplementary Video 1:** 3D rendering of cluster 1 (r1π). https://dx.doi.org/10.6084/m9.figshare.27187599

**Supplementary Video 2:** 3D rendering of cluster 2 (r1π bundle). https://dx.doi.org/10.6084/m9.figshare.27187935

**Supplementary Video 3:** 3D rendering of cluster 3 (small, no axon). https://dx.doi.org/10.6084/m9.figshare.27187941

**Supplementary Video 4:** 3D rendering of cluster 4 (small). https://dx.doi.org/10.6084/m9.figshare.27187944

**Supplementary Video 5:** 3D rendering of cluster 5 (ipsi-lateral/caudal projection). https://dx.doi.org/10.6084/m9.figshare.27187947

**Supplementary Video 6:** 3D rendering of cluster 6 (contralateral/caudal projection). https://dx.doi.org/10.6084/m9.figshare.27187950

**Supplementary Video 7:** 3D rendering of cluster 7 / afferent cluster 1 (ipsilateral, diffused). https://dx.doi.org/10.6084/m9.figshare.27187953

**Supplementary Video 8:** 3D rendering of the afferent cluster 2 (ipsilateral afference). https://dx.doi.org/10.6084/m9.figshare.27187956

**Supplementary Video 9:** 3D rendering of the afferent cluster 3 (contralateral afference). https://dx.doi.org/10.6084/m9.figshare.27187959

**Supplementary Video 10:** 3D rendering of the afferent cluster 4 (vIPN afference). https://dx.doi.org/10.6084/m9.figshare.27187962

**Figure S1:**
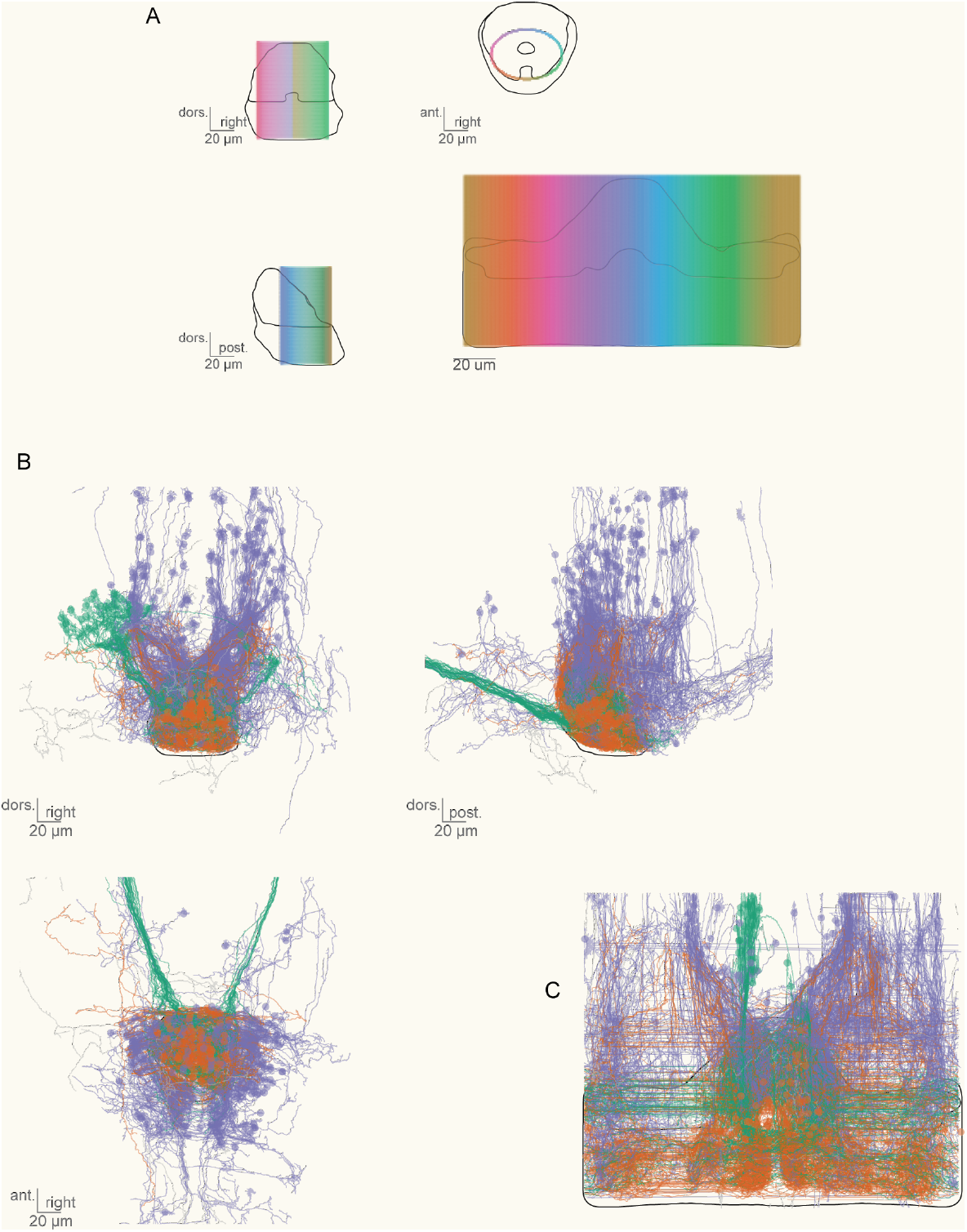
Flattened representation of the IPN. A) Visualization of the ellipse that was used to project the volumetric data in the 2D visualization of the IPN that is used throughout the article (shown in the bottom right corner). B) Visualization of all cells in the cluster, color-coded as in Figure 1: green for habenular neurons, orange for neurons with the soma in the IPN; and purple for neurons with the soma in the dorsal hindbrain.

**Figure S6:**
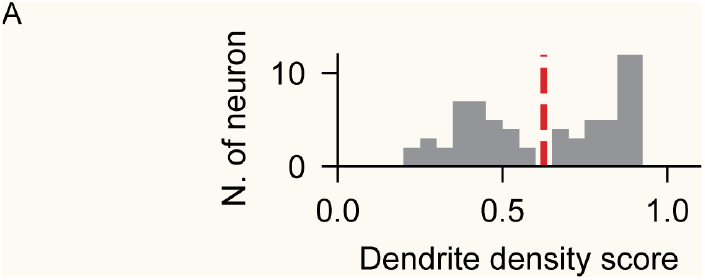
Distribution of dendritic densities in the vIPN. A) Histogram representing the distribution of dendritic densities for all neurons in the vIPN. The density was defined counting the number of other nodes from the same skeletons in a sphere of radius 20 μm. The threshold shows the separation between low and high density neurons as defined in the main text.

**Figure S7:**
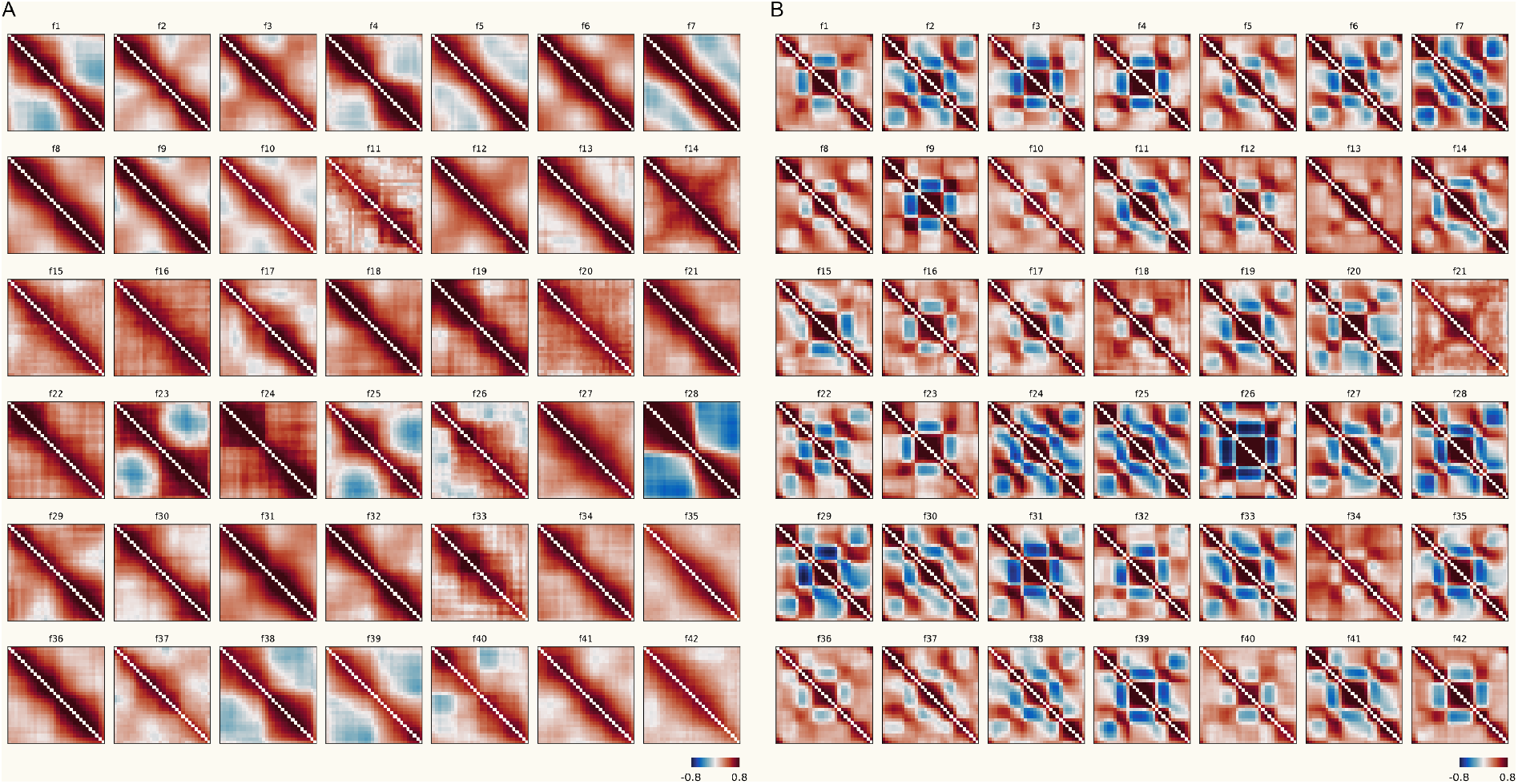
Spontaneous activity in Hb axons. A) Correlation matrix of the spontaneous activity in different medio-lateral positions of the dIPN in each fish. The binning follows Figure 7A. B) Same as A for different ellipsoid positions of the vIPN. The binning follows Figure 7B.

